# Antagonistic regulation of HBZ splicing by hnRNPA1 and hnRNPH1 drives HTLV-1 leukemogenesis

**DOI:** 10.64898/2026.07.20.739521

**Authors:** Julie Tram, Célima Mourouvin, Laetitia Marty, Anika Marie-Delcasse, Arnaud Lecante, Raymond Césaire, Philippe Hélias, Stanie Gaëte, Véronique Baccini, Benoît Barbeau, Norbert Donhause, Andrea K. Thoma-Kress, Jean-Michel Mesnard, Jean-Marie Peloponese

**Affiliations:** Université Montpellier (UM), Montpellier, France; Institut de Recherche en Infectiologie de Montpellier (IRIM), CNRS, Montpellier, France; PCCEI Inserm - Université des Antilles, Pointe-à-Pitre, France; Département de Radiothérapie-Oncologie-Hématologie, Centre Hospitalier Universitaire de La Guadeloupe, Pointe-à-Pitres, France; Karubiotec TM, CRB du CHU de la Guadeloupe, Pointe-à-Pitre, France; Laboratoire d’Hématologie CHU de la Guadeloupe Pointe à Pitre Guadeloupe; Department of Biological Sciences, Université du Québec à Montréal, Montréal, Canada; Harald zur Hausen Institute of Virology, Uniklinikum Erlangen, Friedrich-Alexander-Universität Erlangen-Nürnberg (FAU), Erlangen, Germany

**Keywords:** Adult T-cell leukemia, alternative splicing, hbz, hnRNPA1, hnRNPH1

## Abstract

Adult T-cell leukemia/lymphoma (ATL) is a highly aggressive leukemia driven by Human T-cell Leukemia Virus type 1 (HTLV-1) and remains largely refractory to current therapies. Although *hbz* is the only viral transcript consistently expressed in acute ATL, the extent to which its alternative splicing shapes disease biology remains unknown. Here, we demonstrate that the splicing of *hbz* plays a key role in driving cancer development in ATL. Quantitative analyses in HTLV-1-infected cell lines and primary samples revealed a striking enrichment of the spliced isoform HBZ_SP1 (over 200-fold) in CD4⁺ T cells from ATL patients, whereas the unspliced transcript (usHBZ) predominates in CD8⁺ T cells. Despite robust transcription, the usHBZ protein was undetectable, whereas HBZ_SP1 accumulated rapidly, identifying it as the main isoform in CD4⁺ T cells from ATL patients. Furthermore, only HBZ_SP1 drove cellular transformation and conferred marked resistance to chemotherapeutic stress. Mechanistically, we identify a splicing regulatory axis centered on hnRNPA1 and hnRNPH1. Both proteins bind *hbz* pre-mRNA, but exert opposing effects: hnRNPA1 represses splicing, whereas hnRNPH1 promotes production of the oncogenic HBZ_SP1 isoform. Perturbation of this balance reprograms HBZ isoform expression and alters leukemic cell fitness. Collectively, our findings establish that HBZ inhibits hnRNPA1 transcription, therefore allowing HTLV-1 to hijack host RNA splicing and to generate an oncogenic isoform that drives transformation and chemoresistance. These results uncover a previously unrecognized post-transcriptional mechanism of viral leukemogenesis and position HBZ splicing and its regulators as therapeutic targets in ATL.

**Graphical abstract:** 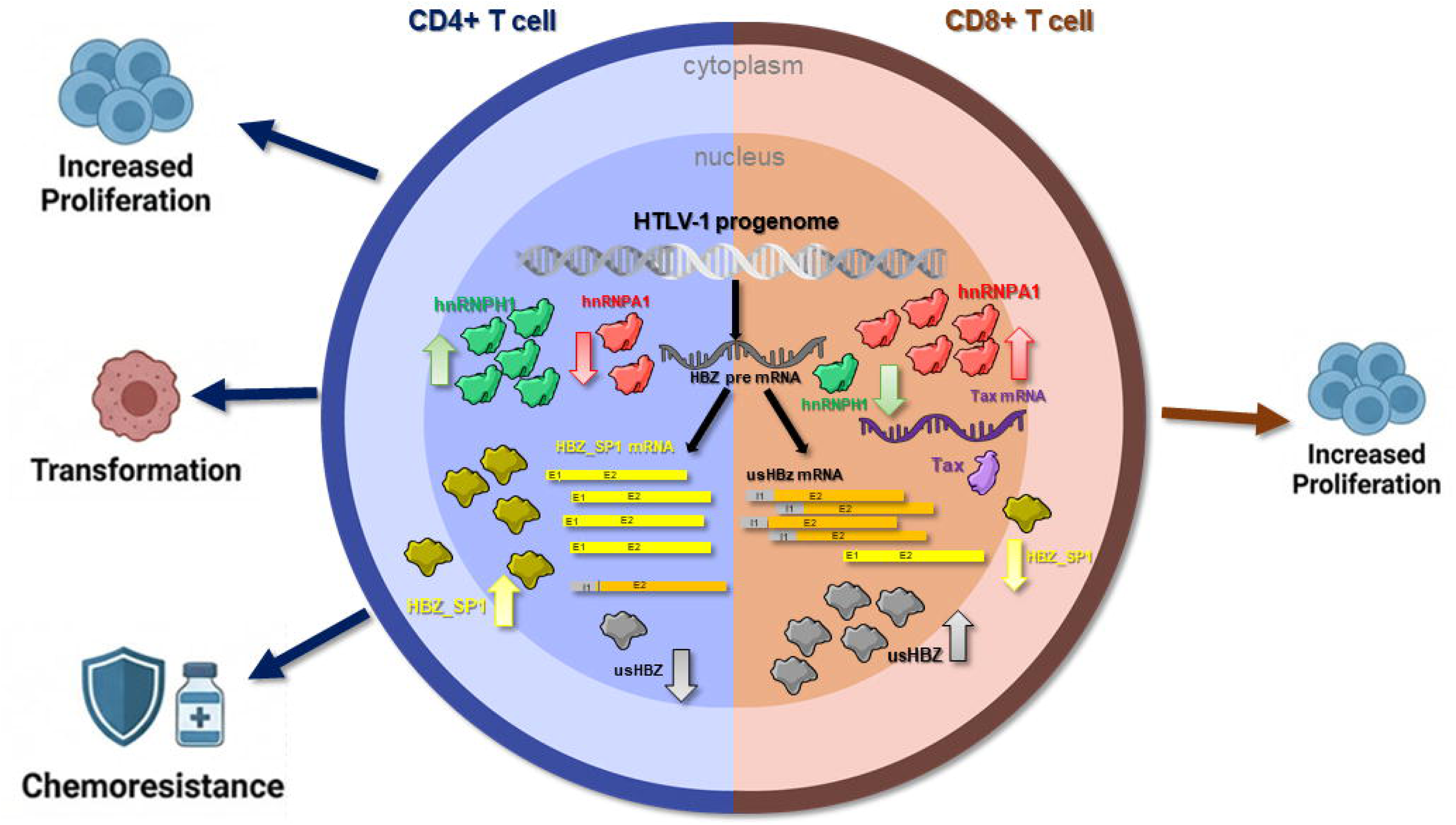

**Mechanistic model of HBZ splicing regulation.** This simplified graphical abstract summarizes the highlights of our study. Here, we hypothesize that in CD4^+^ T cells infected by HTLV-1, transcription and splicing reprogramming lead to a preferential expression of the HBZ_SP1 oncogenic isoform, specifically due to an overexpression of the splicing activator hnRNP H1and repression of hnRNP A1 by HBZ itself via C/EBPα. HBZ_SP1 is consistently expressed and then drives cell transformation and chemoresistance. In contrast, in CD8^+^ T cells infected with HTLV-1, the HBZ-mediated downregulation of hnRNP A1 is overcome, leading to inhibition of HBZ splicing and increased expression of the less oncogenic isoform usHBZ.

## INTRODUCTION

Chemoresistance remains a major obstacle to effective cancer treatment and can arise through intrinsic mechanisms or be acquired following drug exposure^1–6^. Multiple mechanisms contribute to chemoresistance, including increased drug efflux via ATP-binding cassette transporters, enhanced DNA repair, evasion of apoptosis, and changes in the tumor microenvironment^1–8^. Growing evidence now identifies alternative RNA splicing as a critical contributor to tumor progression and chemoresistance, enabling the generation of oncogenic isoforms that promote survival and adaptation^2,5,9,10^.

RNA splicing is a crucial post-transcriptional process ^11–13^ controlled by the spliceosome and its associated factors, including the serine-rich (SR) and heterogeneous ribonucleoprotein (hnRNP) families ^14–17^. Dysregulated splicing is increasingly recognized as a hallmark of cancer^18–21^ and is frequently driven by mutation or altered expression of splicing factors such as SF3B1, SRSF2, U2AF1, and ZRSR2^20,22,23^. These mutations can cause exon skipping, intron retention, and the creation of novel oncogenic splice variants. Splicing errors may also affect genes involved in apoptosis, cell cycle regulation, and DNA repair^20,22,23^, such as the anti-apoptotic BCL-X and MCL1 isoforms which can promote the survival of cancer cells ^24^. Therefore, targeting splicing therapeutically, either by correcting faulty splicing or by exploiting cancer-specific splice variants, offers a promising innovative approach in oncology^15,18,19,25,26^

Adult T-cell Leukemia/Lymphoma (ATL) is an aggressive hematologic malignancy that affects CD4^+^ T cells and is caused by infection with the Human T-Leukemia Virus type 1 (HTLV-1)^27–34^. ATL is characterized by genomic instability, epigenetic changes, and immune evasion^27–34^. ATL has a very poor diagnosis, with a median survival time of less than one year for the acute and lymphoma clinical subtypes^35^. While the viral protein Tax promotes early transformation, it is frequently silenced in established disease due to immune pressure^27–34^. Conversely, HTLV-1 basic leucine zipper factor (HBZ) is consistently expressed in ATL cells and is essential for maintaining the malignant phenotype^27–34,36^.

HBZ is transcribed from the minus strand of the HTLV-1 provirus and exists in both RNA and protein forms^32,37–41^. *hbz* mRNA promotes T-cell proliferation, whereas the protein form modulates transcription factors such as NF-κB, AP-1, and CREB to support cell survival, immune evasion, and proliferation^32,37–41^. Additionally, HBZ enhances Treg-like phenotypes and impairs anti-viral immune responses by modulating FOXP3 expression and function^32,37–41^. Emerging evidence suggests that HBZ interacts with RNA-processing factors and may influence splicing^42,43^. However, the role of HBZ alternative splicing in ATL pathogenesis remains poorly understood. Here, we demonstrate that the HBZ splicing ratio is inverted between CD4^+^ and CD8^+^ T cells isolated from patients with ATL. To understand the mechanisms regulating *hbz* splicing, we profiled over 20 splicing factors (SFs) with predicted binding sites within *hbz* pre-mRNA, and identified a regulatory axis involving hnRNPA1 and hnRNPH1 that controls isoform balance and links viral RNA processing to chemoresistance. Our findings highlight key regulators of HBZ isoform splicing, offering promising new avenues for ATL treatment. Targeting pathways involved in HBZ splicing could enhance chemosensitivity and improve patient outcomes, addressing the urgent need for effective therapies in this aggressive disease.

## MATERIALS AND METHODS

### HTLV-1 patients’ samples

Samples include non-infected individuals (NI); HTLV-1 infected patients without declared malignancy (AC for asymptomatic carriers); patients with tropical spastic paraparesis/HTLV-1 associated myelopathy (HAM/TSP), and patients with acute ATL (ATL). These samples were collected at the Teaching Hospital (CHU) of Martinique and the CHU of Guadeloupe (both in the French West Indies). The diagnosis of ATL was based on clinical features, hematological findings, presence of the HTLV-1 provirus in leukemic cells, and detection of anti-HTLV-1 antibodies in patients’ sera. ATL was subtyped according to the JLSG criteria^44^. Peripheral blood samples were used in accordance with French bioethics laws regarding the use of biological samples. The Comité de Protection des Personnes approved all experiments involving patient samples. Serum and blood samples were obtained from the processing of biological samples by the Biological resources center of Martinique (CeRBiM),, and the Biological resources center of Guadeloupe (KaruBioTech).

### Constructs

pcDNA3.1 usHBZ-Myc, pcDNA3.1 HBZ_SP1-Myc, pcDNA3.1 HBZ-SM-Myc, pcDNA3.1 HBZΔ2ATG-Myc, pcDNA3.1 HBZ(LLXAA)2-Myc, pcDNA3.1 HBZ-ΔZip Myc, K30-asLuc and Hpx-Tax were previously described^45^. RG6 and Flag-MBNL3 were kindly gifted by Thomas Cooper (Addgene plasmids #80167 and #96901)^46^. CMV-LUC2CP/intron/ARE (Luc-i) and CMV-LUC2CP/ARE (Luc) were generously donated by Gideon Dreyfuss (Addgene plasmids #62858 and #62857)^47^. p3x-FLAG-hnRNPH1 was a gift from Mariano Garcia-Blanco (Addgene plasmid #21925). pcDNA-Flag-hnRNPA1 and pETR3 were purchased from Genscript. pCMV-6-CEBPα (human) was purchased from OriGene, and pLightSwitch-hnRNPA1 promoter from Active Motif. MISSION® pLKO.1-puro shRNA hnRNPA1 was purchased from Sigma-Aldrich.

### Primary CD4 and CD8 T lymphocytes purification

PBMCs were isolated from EDTA-anticoagulated blood samples using the Ficoll density gradient method. CD8^+^ and CD4^+^ T-lymphocytes were sorted with the EasySep™ Human CD8 Positive Selection Kit II and the EasySep™ Human CD4^+^ T Cell Isolation (STEMCELL Technologies). CD8^+^ and CD4^+^ T-lymphocytes from ATL patients (∼5×10^5^ cells/mL) were then resuspended in 2 mL of ImmunoCult™-XF T Cell Expansion Medium, supplemented with ImmunoCult™ Human CD3/CD28/CD2 T Cell Activator and 10 ng/mL of RhIL-2 (STEMCELL Technologies). When appropriate, valproate (2-n-propylpentanoic acid, VPA) was added to the medium at a concentration of 5 mM. Cells and culture supernatants were harvested after various incubation times at 37°C in 5% CO_2_, from 0 (D0) up to 5 days (D5), depending on the analysis.

### Cell culture and transfection

CD4^+^ T-lymphocytes, human-derived T-cell lines Jurkat (ATCC® TIB-152™), HuT78 (ATCC® TIB-161™), the HTLV-1 transformed cell lines, HuT-102 (ATCC® TIB-162™), C8166 (ECACC 88051601), and ATL-2 (CVCL_A6TF), JuanaW^48^, were propagated in RPMI 1640 supplemented with 10% of fetal calf serum (FCS), 20U/ml rhIL-2 (STEMCELL Technologies) when necessary, and transfected by electroporation using a GenePulser X (Bio-Rad) according to the manufacturer’s protocol. NIH3T3 (ATCC® CRL-1658™), Cos7 (ATCC® CRL-1651™), and HEK 293 cell lines (ATCC® CRL-1573) were propagated in DMEM supplemented with 10% FCS and 100U/ml penicillin/streptomycin and transfected according to the manufacturer’s protocol using Pei Max (Polyplus Transfection).

### RNA extraction and RT-qPCR analysis

Total RNA was extracted from whole cells using TriReagent (Invitrogen), following previously described protocols^49^. After performing reverse transcription with the 5x master mix (ABM), transcript levels were quantified via qPCR using SYBR Green PCR Master Mix (Roche Diagnostics). Gene-specific primers used for qPCR are described in Supplementary Table 1. The qPCR data were analyzed using LightCycler 480 Software (Roche Diagnostics).

### Western-blotting

Primary cells, stable cell lines, T-cell lines, or transfected cells were lysed in RIPA buffer containing 10 mM Tris–HCl (pH 7.4), 150 mM NaCl, 1% NP-40, 1 mM EDTA, 0.1% SDS, and 1 mM DTT. Cell lysates were further disrupted by sonication. Samples were separated on SDS-PAGE gels (Bio-Rad) and transferred using the Turbo RTA Transfer Kit (Bio-Rad). Primary antibodies used to probe blots were listed in Supplementary Table 2. HRP-conjugated secondary antibodies targeting mouse or rabbit IgG (Abcam) were utilized for detection. Protein bands were visualized through enhanced chemiluminescence using a Chemidoc Imaging System (Bio-Rad). When necessary, band intensities were quantified using ImageJ and normalized to the loading control protein levels.

### Proliferation and Chemoresistance Assays

HBZ-expressing cells were exposed for 48 hours to Etoposide (Sigma-Aldrich) or DMSO as a negative control. Cell viability was assessed using the PrestoBlue viability reagent (Thermo Fisher) by measuring fluorescence (excitation at 560 nm; emission at 600 nm) according to the manufacturer’s instructions and cell survival was normalized to untreated controls.

### Transformation assay (soft agar)

Soft agar in 96-well plate assays was prepared as described by Ke et al.^50^ The cells were cultured in a humidified 37°C incubator with 5% CO_2_ for 1-2 weeks. Colony growth and cell viability were measured using Prestoblue (Invitrogen) by reading the OD at 570nm on a SPARK 10M microplate reader (TECAN). Colony pictures were taken using an Evos microscope (Nikon).

### RNA immunoprecipitation assay

HEK293 cells were co-transfected with the molecular clone K30-3’-4089^51^ and the Flag-hnRNPA1 or with Flag-hnRNPH1 plasmids. 48 hours after transfection, cells were collected using cold CRB buffer (40 mM Tris, pH 7.5, 1 mM EDTA, 150 mM NaCl). RNA immunoprecipitations were performed using the Magna RIP™ RNA-Binding Protein Immunoprecipitation Kit (Merck Millipore), according to the manufacturer’s protocol. After washing, the immunoprecipitated complexes were analyzed by SDS-PAGE and used for RNA extraction and quantification by RT-qPCR as described previously.

### Minigenes splicing assay

To assess the impact of HBZ on splicing, three minigenes were used: the Luc-i-Luc minigene^47^, the RG6 minigene^46^, and the K30’-as-Luc minigene^51^. Briefly, HEK293 or HuT78 cells were co-transfected with one of these minigenes and either HPX-Tax, pcDNA3.1 usHBZ-Myc, or pcDNA3.1 HBZ_SP1-Myc. The impact of HBZ on splicing was measured by luminescence for Luc-i-Luc and K30’-as-Luc minigenes, and by GFP and dsRed fluorescence for the RG6 minigene, using a SPARK 10M microplate reader (TECAN). The effect of HBZ on transcription was assessed by measuring *firefly* luciferase mRNA levels by RT-qPCR.

### Luciferase assay

To measure luciferase activity, HEK cells were transfected with a DNA mixture containing 100ng of the LightSwitch hnRNPA1 promoter and 100ng of the pAct-β-galactosidase construct. After transfection, cells were collected with cold CRB buffer, rinsed with PBS, and lysed with Passive Lysis Buffer (Promega). The LightSwitch luciferase reporter assay system reagents (Active Motif) were used for luciferase detection, and the Galacto-Star kit (Tropix) for β-galactosidase assays, following the manufacturers’ protocols. Luciferase and β-galactosidase activities were measured using a SPARK 10M microplate reader (TECAN). Luciferase signals were normalized to β-galactosidase activity to account for transfection efficiency.

### Statistical analysis

One-way ANOVA or two-tailed unpaired Student’s t test was used for *in vitro* cell line and primary cell experiments, including luciferase assay, minigene assays, RT-qPCR, RIP assay, and cell growth assay. Error bars show SD. Differences were considered significant at *p<0.1, **p<0.01, ***p<0.001 and ****p<0.0001.

## RESULTS

### HBZ isoform expression shifts toward the spliced form 1 during ATL progression

To determine whether HBZ RNA splicing is altered during disease progression, we quantified usHBZ and HBZ_SP1 transcripts in CD8^+^-depleted PBMCs from asymptomatic carriers (AC), patients with HTLV-1-associated Myelopathy/tropical spastic paraparesis (TSP), and patients with Adult-T-cell Leukemia (ATL). While usHBZ levels showed only modest variation across the different groups (Figure 1A), HBZ_SP1 transcripts were increased in ATL samples (Figure 1B). This difference became more apparent when examining the HBZ_SP1/usHBZ ratio, which is higher in ATL patients (median >361) than in AC (∼33) (Figure 1C). Using our monoclonal anti-HBZ antibody (AB49), we observed that HBZ_SP1 protein accumulates over time in *ex vivo*-cultured TSP- and ATL-derived lymphocytes (Figure 1D–E). In contrast, despite detectable usHBZ mRNA and confirmed antibody reactivity toward the usHBZ protein (Sup_Figure S1), no corresponding protein signal was observed in the same samples. Consistent with these observations, HTLV-1-infected cell lines also displayed preferential expression of HBZ_SP1, as measured by RT-qPCR and further highlighted by the ratio (Sup_Figure S1B–D), reinforcing the idea that splicing regulation of HBZ is a hallmark of the leukemic state.

**Figure 1.**
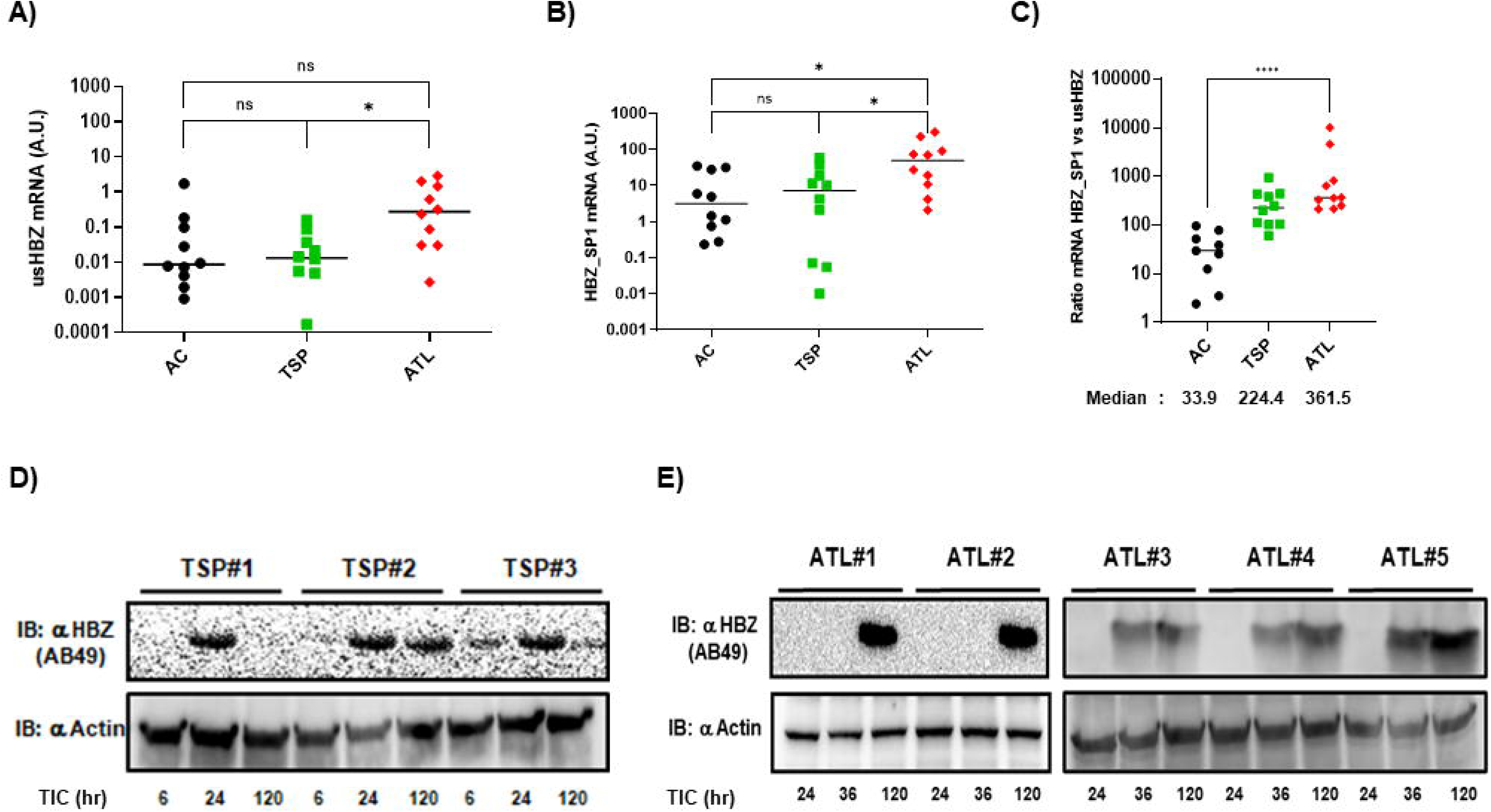
HBZ isoforms expression shifts in ATL. **(A-B)** HBZ isoforms RNA expression was quantified in CD8^+^-depleted PBMCs isolated from 10 HTLV-1-infected asymptomatic carriers (AC), 10 patients with HAM/TSP (TSP), and 10 ATL patients (ATL) by RT-qPCR using isoform-specific primer. **(C)** HBZ_SP1/usHBZ expression ratio was calculated, and the median is shown. Both isoforms are expressed at significantly higher levels in ATL patients compared to control or TSP patients. **(D-E)** Western blot analysis of HBZ protein levels was performed in whole-cell lysates from three HAM/TSP and five ATL patients using the monoclonal anti-HBZ antibody (AB49 clone) with an anti-actin blot as a loading control. HBZ is detectable in all TSP and ATL patients (chronically infected with HTLV-1) starting at 24 hours in TSP patients and 36 hours in ATL patients, with a peak at 5 days (120 hours). (TIC: Time in Culture)

### HBZ spliced isoform 1 drives proliferation, transformation, and chemoresistance

To assess the functional impact of HBZ isoforms, we stably expressed usHBZ or HBZ_SP1 in immortalized murine NIH3T3 cells. RT-qPCR confirmed robust expression of both constructs (Figure 2A), and proliferation assays showed that both isoforms increased cell growth compared with mock cells. However, HBZ_SP1 induced a stronger and sustained proliferative response (Figure 2B). We next evaluated the transforming capacity of each HBZ isoform using a soft-agar assay. Control and usHBZ cells formed few or no colonies, whereas HBZ_SP1 led to increased anchorage-independent growth, in both colony number and size (Figure 2C). The viability of such colonies was assessed, revealing that only HBZ_SP1-expressing cells are continuously proliferating in the soft-agar (Figure 2D). Next, to explore the link between HBZ and drug resistance, we exposed the stable cells to increasing concentrations of etoposide, a chemotherapeutic agent. Cells expressing HBZ_SP1 showed chemoresistance, with an IC50 of ∼100 µM, exceeding the FDA range (∼33 µM)^52^, while usHBZ cells were as sensitive as the mock cells (Figure 2E). These results indicate that the predominant HBZ_SP1 isoform confers key oncogenic properties observed in ATL.

**Figure 2.**
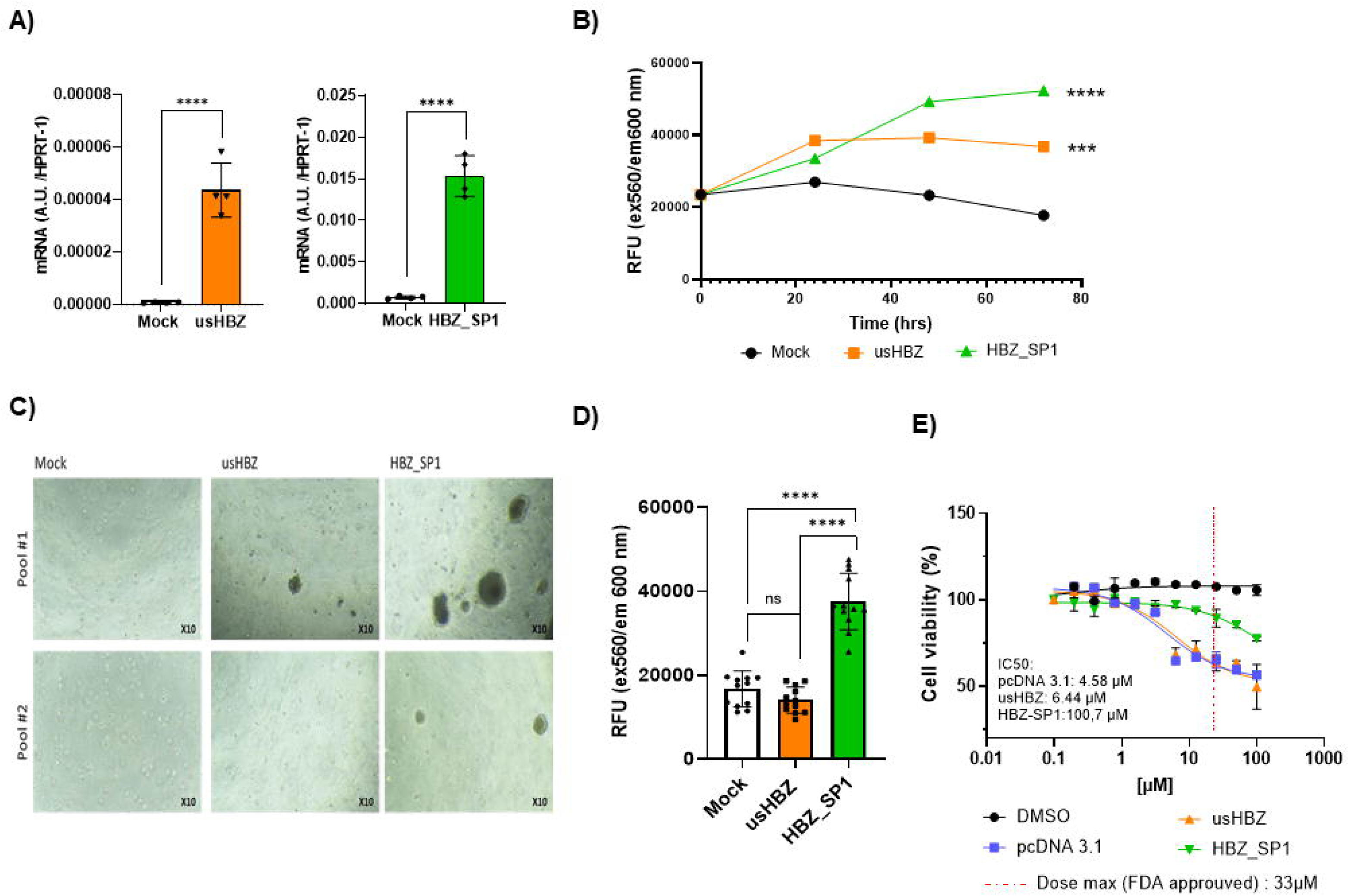
HBZ_SP1 drives cell transformation and drug resistance. **(A)** NIH3T3 cell lines stably expressing either usHBZ or HBZ_SP1 were generated, and the expression level of HBZ isoforms was confirmed by RT-qPCR and compared to the mock cell line represented by NIH3T3 transfected with pcDNA3.1. **(B)** Stable cell lines proliferation under low growth factor conditions (2% FCS) was monitored via fluorescence measures for 72 hours using PrestoBlue viability reagent. Data show HBZ expressing cells survive better than mock cells with HBZ_SP1 cells, even proliferating over time. (**C-D**) Anchorage-independent growth with low FBS was also assessed using a soft-agar assay for 14 days, and cell colony formation was observed under a bright-field microscope, with cell viability assessed using the same method as previously. Representative images for each cell line and 2 independent experiments are shown in **(C),** and viability in **(D)**. HBZ_SP1-expressing cells form more and larger colonies and are viable and growing even after 2 weeks. **(E)** Stable cell lines were treated with increasing concentrations of etoposide or DMSO as a control, and cell viability was assessed using PrestoBlue. HBZ_SP1 cells survive etoposide treatment better than mock and usHBZ cells, even at concentrations exceeding the Dose max indicated by the red dotted line.

### HBZ_SP1 influences pre-mRNA splicing via its activation domain and by inhibiting intron exclusion and promoting exon inclusion

Given the strong enrichment of HBZ_SP1 in ATL, we then investigated whether HBZ could influence RNA splicing and thus its own splicing. For that, we used the Luc-I/Luc minigene developed by Younis et al.^47^. In this assay, if the intron (Luc-I) is included via an alternative splicing event, stop codons prevent luciferase expression (Figure 3A). Co-expressing the minigene with Tax or HBZ_SP1, we observed that HBZ significantly reduced Luc-I splicing efficiency, as indicated by decreased luciferase activity, whereas Tax had no effect (Figure 3B). Co-expression with the Luc minigene confirms that changes in luciferase are due to splicing modulation. We then tested whether HBZ’s effect was proportional to its expression by increasing HBZ levels. The results indeed showed a dose-dependent effect (Figure 3C). Finally, we co-expressed the minigene with either usHBZ or HBZ_SP1. Only HBZ_SP1 reduced splicing while usHBZ had no effect (Figure 3D).

**Figure 3.**
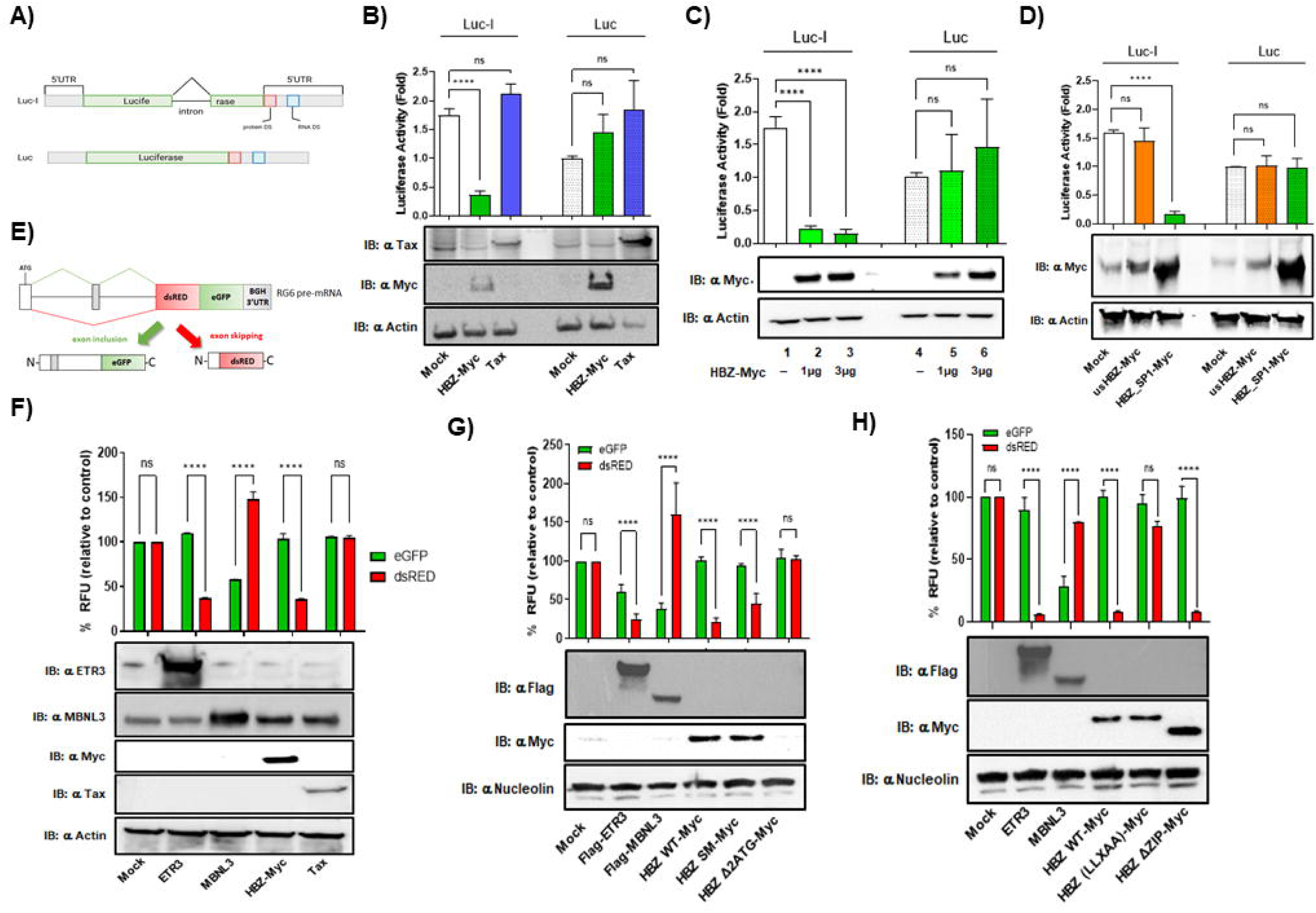
HBZ modulates RNA splicing by promoting exon inclusion. **(A)** Schematic of the Luc-I/Luc minigene, a quantitative reporter system consisting of two luciferase-based minigenes. This Luc-I/Luc minigene has been engineered to express firefly luciferase (F-Luc), with the inserted intron (Luc-I) removed by a constitutive splicing event^47^. . HEK293 cells were co-transfected with either the Luc-I or the Luc minigene and **(B)** either Tax or **(C)** HBZ-Myc, (C) increasing concentration of HBZ-Myc, or **(D)** either HBZ_SP1-Myc or usHBZ-Myc. Luciferase activity was monitored 48 hours after transfection and normalized to actin for control. Results indicate that the RNA splicing inhibition seen for HBZ-Myc can be attributed to the HBZ_SP1 isoform. **(E)** Schematic representation of the RG6 minigene, a dual-fluorescence construct in which inclusion of an alternative exon drives eGFP expression, whereas exon skipping leads to dsRed expression, allowing quantitative assessment of splicing decisions. The RG6 construct is transfected in HEK293 cells with **(F)** HBZ-Myc or Tax, **(G)** HBZ WT, SM, or Δ2ATG, **(H)** HBZ WT, HBZ (LLXAA), or ΔZIP mutant^45^. Measurements of each fluorophore were performed on a Spark 10M microplate reader (Tecan). Corresponding Western blots confirm transfection efficacy and protein expression in each experiment, using either actin or nucleolin as a loading control. All data were presented as the mean of three independent experiments ± standard deviation (SD). Statistical significance was determined by a 1-way ANOVA, ****=p≤0,0001.

To further explore HBZ’s impact on RNA splicing, we used the RG6 dual-fluorescence splicing reporter system developed by Orengo et al.^46^, a dual-fluorescence construct in which inclusion or skipping of an alternative exon drives, respectively, eGFP or dsRED expression. As in previous results, co-expression of the RG6 minigene with Tax did not affect either marker, whereas HBZ expression reduced dsRED fluorescence intensity (Figures 3F, Sup_Figure S2A). To identify HBZ’s implicated regions, we tested multiple HBZ mutants (Sup_Figure S2B). Similar effects on the RG6 were obtained for usHBZ and HBZ_SP1, further confirming the Luc-I/Luc results (Sup_Figure S2E). Since *hbz* mRNA also has functional roles, we tested the HBZΔ2ATG mutant against HBZ-WT or HBZ-SM (a silent mutation). HBZ-SM inhibited dsRED similarly to HBZ-WT, but HBZ-Δ2ATG had no effect (Figure 3G, Sup_Figure S2C), indicating that only the HBZ protein can modulate splicing. In subsequent experiments, we tested mutants affecting the activation domain (HBZ-LLXAA) or the bZIP domain (HBZΔbZIP). Similar to HBZ-WT, the HBZΔbZIP mutant showed reduced dsRED, suggesting that the bZIP domain isn’t required for splicing modulation. However, mutations in the activation domain abolished the effect seen with HBZ-WT (Figure 3H, Sup_Figure S2D). These data identify the activation domain of the HBZ protein as crucial for the interaction and modulation of the host splicing machinery.

### hnRNPA1 is downregulated in ATL and is an HBZ splicing inhibitor

Despite recent studies^40,41^, significant gaps remain in the understanding of how alternative splicing of HBZ pre-mRNA regulates the balance between the two HBZ isoforms. To investigate the regulatory mechanism for HBZ_SP1, we analyzed putative splicing factor binding sites on *hbz* pre-mRNA using ESEfinder2.0 and Human Splicing Finder from Genomis (Sup_Figure S3). Using RT-qPCR, we examined the expression of over twenty candidate splicing factors in HTLV-1-infected cell lines and patient samples. Results revealed widespread dysregulation among several members of the SR or hnRNP families, suggesting a global remodeling of the splicing environment in ATL cells (Figure 4A-B). We next focused on factors with the most pronounced and consistent dysregulation, notably hnRNPA1, H1, and E1. To assess if these candidates directly regulate HBZ splicing, HEK293 cells were co-transfected with hnRNPA1, H1, or E1 and the K30-3’/4089 HBZ clone, and RNA immunoprecipitation (RIP) assays were performed to confirm their binding to *hbz* pre-mRNA. *hbz* pre-mRNA was detectable in the immunoprecipitated fraction for both hnRNPA1 and H1, confirming their binding (Figure 4D-E).

**Figure 4.**
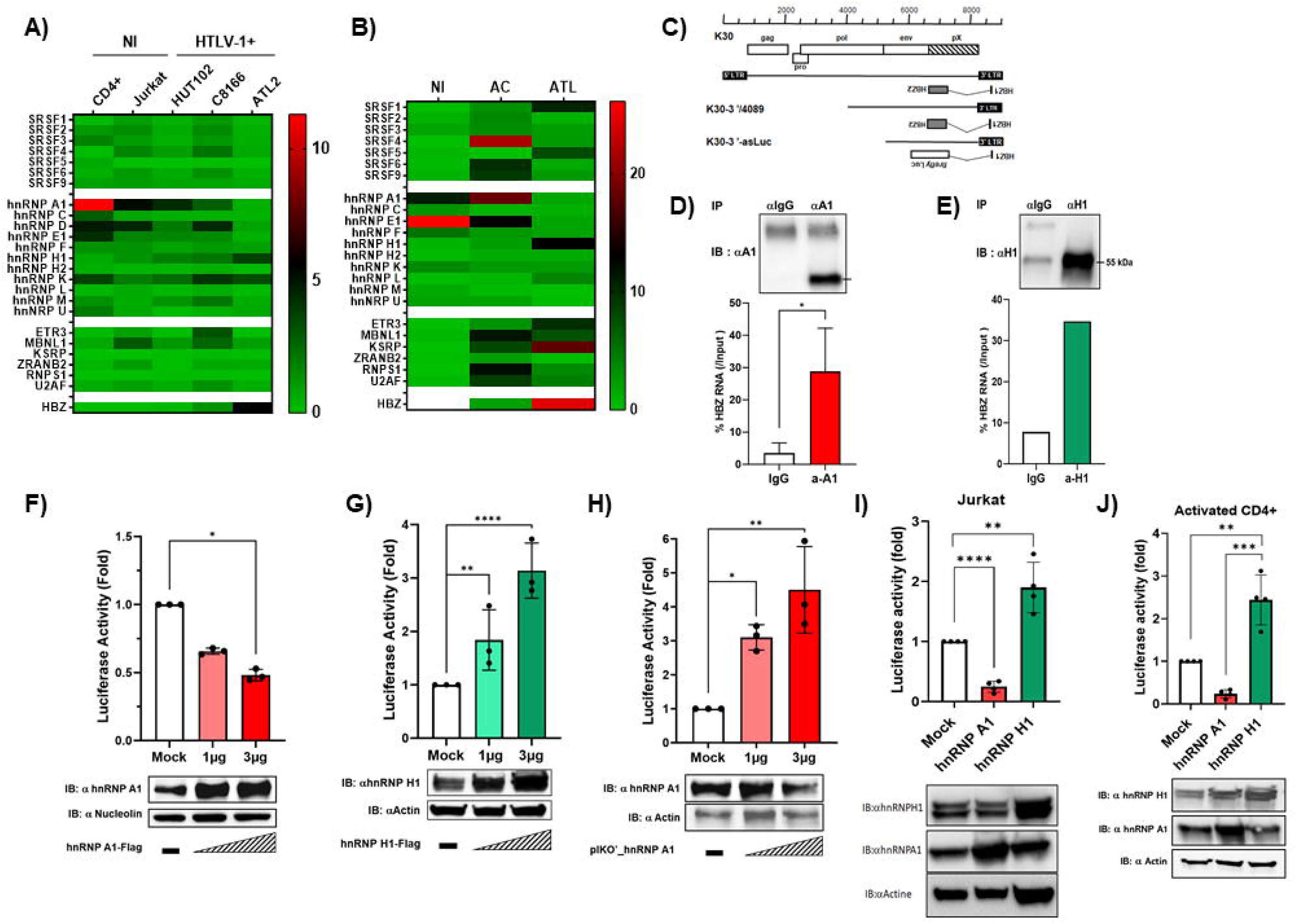
Identification of hnRNP A1 and H1 as HBZ splicing regulators. Heatmap diagrams showing the expression profile of seven slicing factors (SF) from the SRSF family, eleven hnRNPs, six other SF (ETR3, MBNL1, KSRP, ZRANB2, RNPS1, U2AF), and HBZ in **(A)** activated CD4^+^ T lymphocytes, an HTLV-1 negative T cell line (Jurkat) and three HTLV-1 positive T cell lines (HUT102, C81-66 and ATL-2) or in **(B)** CD8^+^-depleted PBMCs from 10 uninfected individuals, 10 HTLV-1 asymptomatic carriers (AC) and 10 patients with acute ATL (ATL), as measured by RT-qPCR using specific primers. **(C)** Schematic diagrams of the K30-3’/4089 proviral DNA construct and the K30-3’-asLuc minigene. The RNA immunoprecipitation (RIP) approach was performed in HEK293 cells co-transfected with (D) hnRNP A1 or (G) H1 and the K30-3’-4089 molecular clone. RNA is immunoprecipitated with hnRNP A1 or H1-specific antibody-coated beads (top panels), and detection of *hbz* pre-mRNA is performed by RT-qPCR using specific primers (lower panels), confirming hnRNP A1 and H1 bind *hbz* pre-mRNA (t-student statistical test, *= p≤0,05). HEK293 cells were transfected with either (F) hnRNP A1, (G) hnRNP H1, or **(H)** shRNA against hnRNP A1, along with the K30-3’-asLuc minigene. *Firefly* luciferase activity was normalized with *Renilla* luciferase activity and both were assessed on a Spark 10M microplate reader (Tecan). Corresponding Western blots confirm the protein expression for each experiment. Overexpression of hnRNP A1 efficiently reduced luciferase activity, whereas shRNA-mediated repression or overexpression of hnRNP H1 had the opposite effect. **(I)** Uninfected T cell line Jurkat and **(J)** activated primary CD4+ T-cells were transfected with either hnRNP A1 or hnRNP H1 along with the K30-3’-asLuc minigene in the same way as described before, and confirmed that hnRNP A1 represses luciferase activity when H1 induces it. Statistical significance was determined by student T-test for RIP and one-way ANOVA with Dunnett’s multiple comparisons post-test for luciferase reporter assays; ns=not significant *= p≤0,05, ** = p≤0,01, ***=p≤0,001, ****=p≤0,0001. All data were presented as the mean of three independent experiments ± standard deviation (SD).

Next, to determine the role of hnRNPA1, H1, and E1 in HBZ splicing, we performed an HBZ minigene assay using the K30-3’-asLuc construct^51^ in which the second exon of *hbz* was replaced with an ATG-less firefly luciferase (F-Luc), thus allowing us to monitor HBZ splicing (Figure 4C). HEK293 cells were co-transfected with K30-3’-asLuc and increasing amounts of hnRNPA1 or H1 plasmids. Overexpression of hnRNPA1 caused a significant decrease in luciferase activity (Figure 4F), without impacting the F-Luc mRNA transcription (Sup_Figure S3A). The same effect can be observed in the Cos7 cells (Sup_Figure S3B), as well as with hnRNPE1 overexpression (Sup_Figure S3C). Conversely, results show that hnRNPH1 promotes *hbz* splicing, highlighting its distinct and opposing role within the hnRNP family (Figure 4G, Sup_Figure S3F). Similarly, siRNA-mediated knockdown of hnRNPA1 also significantly increased luciferase activity and, with it, HBZ splicing (Figure 7D and Sup_Figure S3G). These observations were also confirmed in T-cell models, including Jurkat cells and activated primary CD4⁺ T cells (Figure 4I–J). Together, these results indicate that hnRNPA1 and hnRNPH1 hold opposing functions as a repressor and an activator of HBZ splicing in ATL, respectively.

### HBZ modulates its own splicing by down-regulating hnRNPA1 expression via C/EBPα

To investigate hnRNPA1 regulation in ATL, we generated an HBZ-stably expressing HEK293 cell line. HBZ expression led to a significant decrease in hnRNPA1 mRNA, while other hnRNPs like hnRNPC remained unaffected (Figure 5A–B). The hnRNPA1 protein level is also decreased in the presence of HBZ, as confirmed by Western blot in HEK293 (Figure 5C) and ATL patients’ cells (Figure 5F). Next, we explored transcription factor binding sites on the hnRNPA1 promoter using Ominer from the Signaling Pathways Project. Among the promising candidates, we chose to assess C/EBPα’s role via a luciferase assay, with or without HBZ, using an A1 promoter reporter minigene. We found that HBZ downregulates hnRNP A1 transcription by negating C/EBPα activation of the A1 promoter (Figure 5D). This observed effect was also correlated with a reduction in C/EBPα protein expression in the presence of HBZ (Figure 5E). These observations indicate that HBZ regulates splicing, possibly its own, by indirectly inhibiting hnRNPA1 expression via one of its transcription factors, C/EBPα.

**Figure 5.**
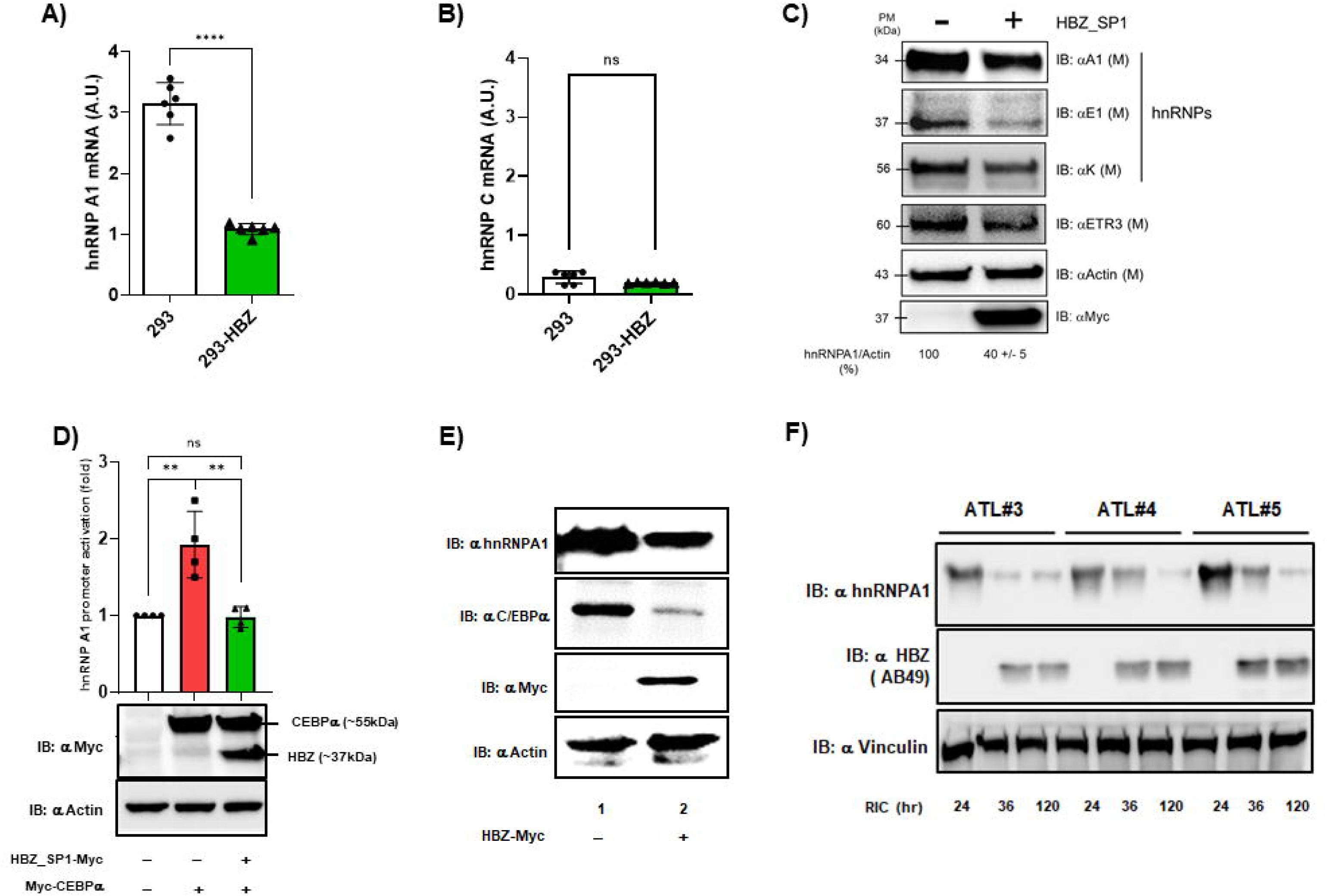
HBZ_SP1 negatively regulated hnRNP A1 expression via C/EBPα. HEK293 cells stably expressing HBZ_SP1 were generated, and expression of hnRNP A1 was monitored by **(AB)** RT-qPCR or **(C)** western blot along with other SF for comparison. Band quantification reveals that hnRNP A1 protein expression is reduced by 60% when HBZ_SP1 is expressed, along with a reduced transcription observed in **(A)**. **(D)** The hnRNP A1 promoter reporter construct was co-expressed with either C/EBPα or HBZ, and promoter activity was assessed via luciferase activation. C/EBPα efficiently activated the hnRNPA1 promoter, while HBZ abrogated the effect when expressed. Corresponding Western blots confirm the protein expression for each experiment. **(E)** hnRNPA1, C/EBPα, and HBZ protein expression were measured in HEK cells stably expressing HBZ_SP1 by western blotting, and reduced expression of both hnRNPA1 and C/EBPα was observed in HBZ-SP1-expressing cells. (**F)** Western blot analysis of ATL patient samples confirmed a consistent reduction in hnRNP A1 expression and its correlation with the presence of HBZ. Statistical significance was determined by Student’s T-test for RT-qPCR and one-way ANOVA with Dunnett’s multiple comparisons post-test for luciferase reporter assays; ns=not significant, *= p≤0,05, ** = p≤0,01. All data were presented as the mean of three independent experiments ± standard deviation (SD).

### HBZ splicing is cell-type dependent and linked to T-cell subsets

CD8^+^ T lymphocytes are infected *in vivo* by HTLV-1 but are not transformed as CD4+ T cells are. Considering our recent findings, we investigated whether HBZ_SP1 is expressed and regulated similarly in CD8^+^ and CD4^+^ T cells. Using RT-qPCR, we assessed the expression of different HBZ isoforms in three CD4^+^ T cell lines chronically infected with HTLV-1 (HuT102, C81-66, ATL-2) and in an HTLV-1-infected CD8^+^ T cell line from an ATL patient, the JuanaW (JW)^48^. Surprisingly, JW cells expressed a high level of Tax transcript (Figure 6A), while HBZ_SP1 mRNA levels were similar in ATL-2 and JW (Figure 6B). usHBZ, on the other hand, was strikingly overexpressed in the CD8^+^ T cell line (Figure 6C), leading the ratio of HBZ_SP1/usHBZ to be completely reversed in the JW cells when compared to CD4^+^ cell lines (Figure 6D). To confirm this, we sorted CD4^+^ and CD8^+^ T cells from four ATL patients and cultured them *ex vivo* for 72 hours in serum-free T cell expansion medium. HBZ_SP1 is strongly enriched in CD4^+^ T cells, whereas Tax and usHBZ predominate in CD8^+^ T cells (Figure 6E-G), with a similar inverted HBZ_SP1/usHBZ ratio (Figure 6H). Quantification of hnRNPA1, E1 and H1 expression in both JW and CD4^+^ and CD8^+^ patient T cells (Sup_Figure S5) confirmed the reversed trend, especially for hnRNPA1, suggesting that the expression and regulation of HBZ_SP1 are cell-specific and linked to the transformation phenotype in ATL.

**Figure 6.**
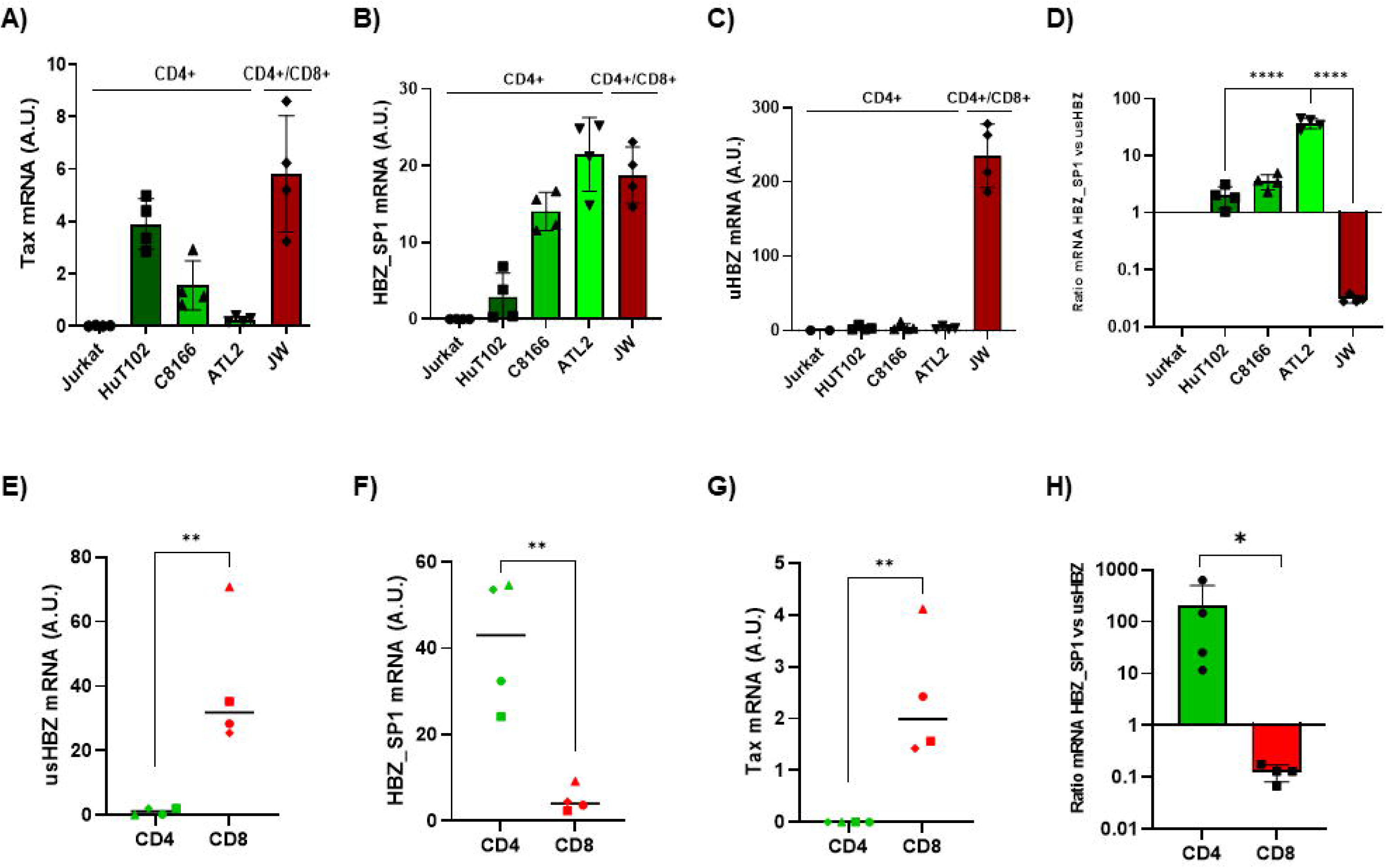
HBZ splicing varies by T-cell subtype in HTLV-1-infected patients. **(A-C)** Tax, usHBZ and HBZ_SP1 isoforms’ respective expressions were measured by RT-qPCR in HTLV-1-uninfected Jurkat T cells, and three HTLV-1-positive CD4+ T cell lines: HuT 102, C81-66, ATL-2, as well as a CD4+/CD8+ HTLV-1-positive T cell line (JuanaW or JW). **(D)** HBZ_SP1/usHBZ expression ratio was measured in T cell lines and plotted. Statistical significance was determined by one-way ANOVA with Dunnett’s multiple comparisons post-test; **** = p≤0,0001. **(E-G)** Expression of usHBZ, HBZ_SP1 isoforms, and Tax was measured by RT-qPCR in CD4^+^ and CD8^+^ T cells isolated from PBMCs of four ATL patients. **(H)** HBZ_SP1/usHBZ expression ratio in CD4^+^ and CD8^+^ T cells was plotted and confirms the reversed ratio in CD8^+^ cells, also visible in the JW, when compared to CD4^+^ or ATL-2 cells. Statistical significance was determined by student T-test; ns=not significant *= p≤0,05, ** = p≤0,01.

### Small molecule targeting of hnRNPA1 alters HBZ isoforms expression balance and rescues ATL cells’ phenotype in CD8^+^ cell types

Finally, we examined whether modulating HBZ splicing via hnRNPA1 could influence ATL cell behavior. To do so, we transfected the T-cell line Hut78 with the K30-3’-asLuc minigene and treated it with 10 µM VPC-80051, an hnRNPA1 inhibitor that disrupts its RNA-binding activity^53^. Western blot analysis confirmed that VPC-80051 treatment did not alter hnRNPA1 expression; however, it significantly increased HBZ splicing (Figure 7A). JW and CD8^+^ T cells from ATL patients were cultivated with or without VPC-80051, and their proliferation was monitored over time. A significant increase in cell growth was observed in both cell types upon treatment with VPC-80051 (Figure 7B-C). This proliferation was also correlated with a significant increase in HBZ_SP1 expression, with no effect on usHBZ or hnRNPA1 transcription (Figure 7D-G). Taken together, our results indicate that small molecule targeting of hnRNPA1 leads to an increase in HBZ_SP1 expression, which in turn promotes cell proliferation through its previously described oncogenic properties.

**Figure 7.**
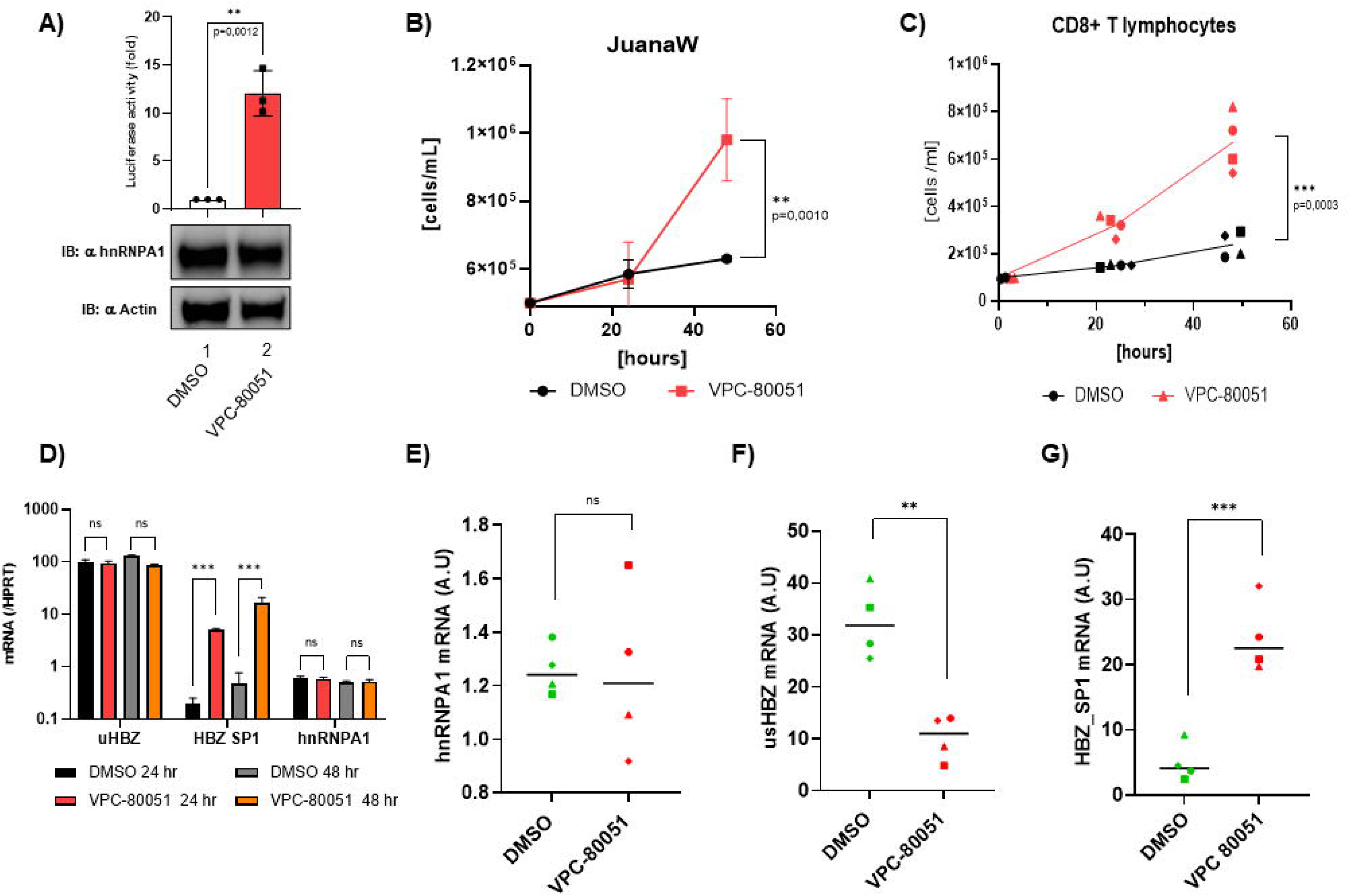
Small-molecule inhibition of hnRNP A1 rescued the ATL phenotype. **(A)** HuT78 cells were transfected with the K30-3’asLuc minigene and treated with 10 µM VPC-80051, an hnRNPA1 inhibitor that disrupts its RNA-binding activity. Firefly luciferase activity was measured and normalized to Renilla luciferase activity, and it increased with VPC treatment. Control Western blots are shown in lower panels. Proliferation curves upon treatment with VPC-80051 for 48 hours were determined by trypan blue exclusion on Countess II Automated Cell Counter (Invitrogen) for **(B)** JW cells and **(C)** CD8^+^ T cells from four ATL patients. Treatment increased cell proliferation in both cases. Statistical significance was determined by two-way repeated-measures ANOVA; ns=not significant, ** = p < 0.001, ***=p≤0,0001. **(D)** JW cells were treated with VPC-80051for 24 or 48 hours, and expression of usHBZ, HBZ_SP1, and hnRNP A1 was measured by RT-qPCR and normalized to HPRT-1 (housekeeping gene). HBZ_SP1 mRNA significantly increased upon treatment. Statistical significance was determined by one-way ANOVA with Dunnett’s multiple comparisons post-test; ns=not significant, *** = p≤0,001. **(E-G)** CD8^+^ T cells from four ATL patients were treated with VPC-80051 for 48 hours, and the expression of usHBZ, HBZ_SP1, and hnRNP A1 was measured by RT-qPCR and normalized to HPRT-1 (housekeeping gene), confirming a switch in expression toward HBZ_SP1. Statistical significance was determined by Student’s t-test; ns=not significant, ** = p≤0,01, ***=p≤0,001.

## DISCUSSION

Our research emphasizes an important aspect of HTLV-1 biology and pathogenesis: the cancer-causing potential of HBZ not only depends on how much is produced but also on how it is spliced and which isoform is expressed and active. Although HBZ expression and biological functions have long been linked to ATL, our results show that a predominance of the HBZ_SP1 isoform acts as a crucial switch in leukemogenesis. Alternative splicing of HBZ produces distinct isoforms with unique biological roles ^51,54^, yet their respective impact on ATL remained underexplored. In this work, we found that the HBZ_SP1 isoform is the main driver of HBZ-mediated cellular transformation and drug resistance mechanisms observed in ATL. Our findings align with the recent report that HBZ_SP1 stabilizes c-Jun and promotes T-cell growth^55^. In contrast, the unspliced isoform (usHBZ), although present at detectable RNA levels, does not appear to express a detectable protein. These results underscore the critical role of post-transcriptional regulation in shaping the expression of the most oncogenic HBZ isoform. Our study also highlights that HTLV-1 hijacks the host’s RNA-processing machinery. RNA-binding proteins and splicing factors are increasingly recognized as key players in cancer biology^43,56^, and our findings further support the role of HBZ in actively reshaping the post-transcriptional regulatory landscape^42^. We identified hnRNPA1 and hnRNPH1 as two regulators of HBZ splicing, whose expression balance creates a regulatory axis that influences the production of the oncogenic HBZ_SP1 isoform. We find here that hnRNP A1 acts as a negative regulator of HBZ splicing, consistent with its well-known role in promoting exon skipping^15,57^. Beyond its role in RNA processing, hnRNPA1 is associated with chemoresistance across various cancers^9,58^, influencing alternative splicing programs that govern how tumors respond to drugs. For instance, hnRNPA1-dependent splicing events can alter sensitivity to cisplatin ^58^ and temozolomide ^9^, and support tumor growth by switching isoforms ^59^. Our data suggest that this pattern may extend to virus-related cancers, indicating that hnRNP A1’s influence on HBZ splicing could help confer chemoresistance as observed in ATL.

In contrast, hnRNPH1 acts as a positive regulator of HBZ splicing, promoting the production of the HBZ_SP1 isoform, consistent with its overexpression in ATL. Recently, hnRNPH1 has been recognized as a significant contributor to tumor progression, especially in glioblastoma, where it governs the splicing of cell cycle-related genes^60^, and in lung adenocarcinoma, where it promotes oncogenic splicing patterns through interactions with signaling pathways ^61^. More broadly, alterations in hnRNPH1 expression contribute to aberrant splicing programs in hematological malignancies^26,62^, reinforcing the relevance of this factor in ATL. Together, our results support a mechanistic model in which the balance between hnRNPA1 and hnRNPH1 is crucial for determining HBZ splicing outcomes. hnRNPA1 restricts HBZ_SP1 production, whereas hnRNPH1 promotes it, creating a regulatory equilibrium that seems to favor the oncogenic isoform expression in ATL. This type of antagonistic regulation is consistent with the broader behavior of hnRNP family members, which often exert opposing effects on splice-site selection, depending on the cellular or pathological context ^16^.

A key aspect of our study is the identification of a feed-forward regulatory loop in which HBZ downregulates hnRNPA1 expression via C/EBPα, thereby preventing hnRNPA1 from inhibiting HBZ splicing in ATL. This dynamic interplay likely fine-tunes viral gene expression across different stages of infection, consistent with recent observations of Tax and splicing-dependent regulation of HBZ expression^63^. Such regulatory circuits may allow HTLV-1 to balance persistence, immune evasion, and oncogenic transformation.

From a translational perspective, our findings highlight a promising potential for therapeutic advancement. The ability to modulate HBZ splicing with the hnRNPA1 inhibitor VPC-80051 supports the idea that targeting RNA-binding proteins can directly enhance leukemic cell vulnerability. With growing interest in targeting splicing factors for cancer treatment^15,26^, hnRNPA1 and hnRNPH1 emerge as promising, possibly complementary, novel candidates. Disrupting the delicate balance between hnRNPA1 and hnRNPH1 could be a key strategy to reduce HBZ_SP1 production, thereby reducing the risk of cell transformation or drug resistance, and paving the way for more effective therapies.

Finally, the findings on how HBZ splicing differs between CD4⁺ and CD8⁺ T cells highlight the crucial role of cellular context in shaping viral oncogenesis. Since ATL arises from CD4⁺ T cells, the predominant accumulation of HBZ_SP1 in this specific cell type suggests that the unique splicing environment within these cells significantly influences disease development. In contrast, infected CD8^+^ T cells, that express lower levels of HBZ_SP1, do not enter an oncogenic pathway. These observations align with previous studies linking HTLV-1 gene expression to chromatin structure and host cell identity, further emphasizing the complex interplay that drives pathogenesis.

In conclusion, our study establishes alternative splicing as a central driver of HTLV-1-associated leukemogenesis, integrating viral gene regulation with host RNA metabolism and chemoresistance. By uncovering a regulatory axis involving hnRNPA1 and hnRNPH1, we provide mechanistic insights into how RNA splicing controls oncogenic output and identify novel, promising vulnerabilities that could be exploited therapeutically in HTLV-1 infection management and ATL treatment.

## Statement on the Use of Generative AI

The Grammarly AI Tool was used to edit the manuscript and improve its clarity. All intellectual content, research design, and data analysis were carried out solely by the authors.

## Authors’ contributions

Peloponese JM designed the research. Tram J., Marty L., Marie-Delkasse A., Belrose G., Donhauser N. & Mourouvin C. performed research. Kress-Thomas A., Barbeau B., Mesnards J.M., Césaire R., Hélias P., and Baccini V. contributed vital new reagents or analytical tools, Peloponese JM and Tram J. performed data analysis and interpretation, designed the figures, wrote and reviewed the manuscript.

## Data Availability Statements

The authors confirm that the data supporting the findings of this study are available within the article and its supplementary materials.

## Acknowledgments

The Fondation pour la Recherche Médicale supported this work (Equipe FRM DEQ20161136701). J. T. was supported by a doctoral grant from the French Ministry of Research and CBS2 doctoral school. C M. was supported by a grant from the Ligue Contre le Cancer, Comité Guadeloupe. L.M. is the recipient of an ANR doctoral fellowship (ANR ANR-22-CE18-0019-01). The authors acknowledge the Centre de Ressources Biologiques de Martinique (CeRBiM), CHU Martinique, France, and de Guadeloupe (KAruBiotech), CHU Guadeloupe, for managing patient samples.

**Sup_Figure 1, related to Figure 1. HBZ isoform expression shifts in HTLV-1 cell lines. (A)** HEK293 cells were transfected with either usHBZ-Myc or HBZ_SP1-Myc, and HBZ protein levels were assessed by western blotting using the monoclonal anti-HBZ (AB49 clone). **(B-C)** HBZ isoforms’ expression was quantified in HTLV-1 negative T cell line Jurkat and in HTLV-1 positive T cell lines HuT102, C81-66, and ATL-2 using RT-qPCR with specific primers, and **(D)** HBZ_SP1/usHBZ expression ratio is calculated. Statistical significance was determined by Student’s t-test; * = p≤0,05

**Sup_Figure 2, related to Figure 3. RG6 minigene fluorescence. (A)** HEK293 cells were transfected with the RG6 minigene and either pcDNA 3.1 (Mock), HBZ_SP1-Myc WT, or HpX Tax. Representative images of EGFP and dsRED observed under epifluorescence microscopy. **(B)** Different HBZ deletion mutants were generated and tested to identify which HBZ domain is responsible for the splicing alteration, including a silent mutation (SM), a protein-deficient (Δ2ATG), an activation domain (LLXAA), and a bZIP (ΔZIP) mutant. **(C)** Transfection of the RG6 construct in HEK293 cells with HBZ-WT, HBZ-SM or HBZΔ2ATG. The remaining tested mutants are shown in **(D)**. Measurements of each fluorophore were performed on a Spark 10M microplate reader (Tecan). Representative images are shown and relative fluorescence is plotted in **(E)**. Corresponding Western blots confirm the efficacy of transfection and protein expression for each experiment. All data were presented as the mean of three independent experiments ± standard deviation (SD). Statistical significance was determined by a 1-way ANOVA.

**Sup_Figure 3, related to Figure 4. Identification of hnRNP A1 and H1 as HBZ splicing regulators.** Bioinformatic analysis of splicing factors binding sites around the donor and acceptor site of HBZ was performed using ESEfinder2.0 (https://esefinder.ahc.umn.edu/tools/ESE2/) and the Human Splicing Finder from Genomis (https://www.genomnis.com/). The nucleotide sequences around the donor **(A)** and acceptor **(B)** sites of HBZ were analyzed to identify putative splicing factor binding sites. **(C)** The K30-3’-asLuc minigene was co-transfected with increasing amounts of hnRNP A1 in either HEK293 or Cos7 cells. **(C)** After 48 hours, firefly luciferase mRNA was quantified by RT-qPCR to ensure the luciferase activity readout is not impacted by the luciferase transcription. **(D)** Splicing efficiency was measured at 48 hours by Firefly luciferase activity, normalized to Renilla luciferase activity. Cell extracts were then analyzed by Western blotting with anti-hnRNP A1 and anti-actin antibodies. (**E)** HEK293 cells were transfected with K30-3’-asLuc minigene and increasing concentration of hnRNP E1 and splicing was assessed as previously described via luciferase activity measure. Cell extracts were then analyzed by Western blotting with anti-hnRNPA1 and anti-actin antibodies. RT-qPCR control experiment for HEK293 cells transfected with K30-3’-asLuc minigene and **(F)** dose-dependent overexpression of hnRNP H1, or **(G)**increasing amounts of shRNA against hnRNP A1 to ensure the luciferase activity readout is not impacted by the luciferase transcription. Statistical significance was determined by Student’s T-test for RT-qPCR and one-way ANOVA with Dunnett’s multiple comparisons post-test for luciferase reporter assays; ns=not significant, ** = p≤0,01, ***=p≤0,001. All data are presented as the mean of three independent experiments ± standard deviation (SD).

**Sup_Figure 5, related to Figure 6. hnRNPA1 is overexpressed in CD8^+^ T lymphocytes from ATL patients.** hnRNP A1, E1 and H1 expression was quantified by RT-qPCR in **(A-C)** CD4^+^ T cell lines: HTLV-1-negative Jurkat cells, and three HTLV-1-positive T cell lines: HuT 102, C81-66, ATL-2, and one CD4^+^/CD8^+^ HTLV-1-positive T cell line JuanaW (JW) (D-F); or **(D-F)** in CD4^+^ and CD8^+^ T cells isolated from PBMCs of four ATL patients. Statistical significance was determined by Student’s t-test; ns=not significant, *= p≤0,05, ** = p≤0,01, ****=p≤0,0001.

