## Supplementary figures and images for "Antagonistic regulation of HBZ splicing by hnRNPA1 and hnRNPH1 drives HTLV-1 leukemogenesis"

### supplementary Figure 1

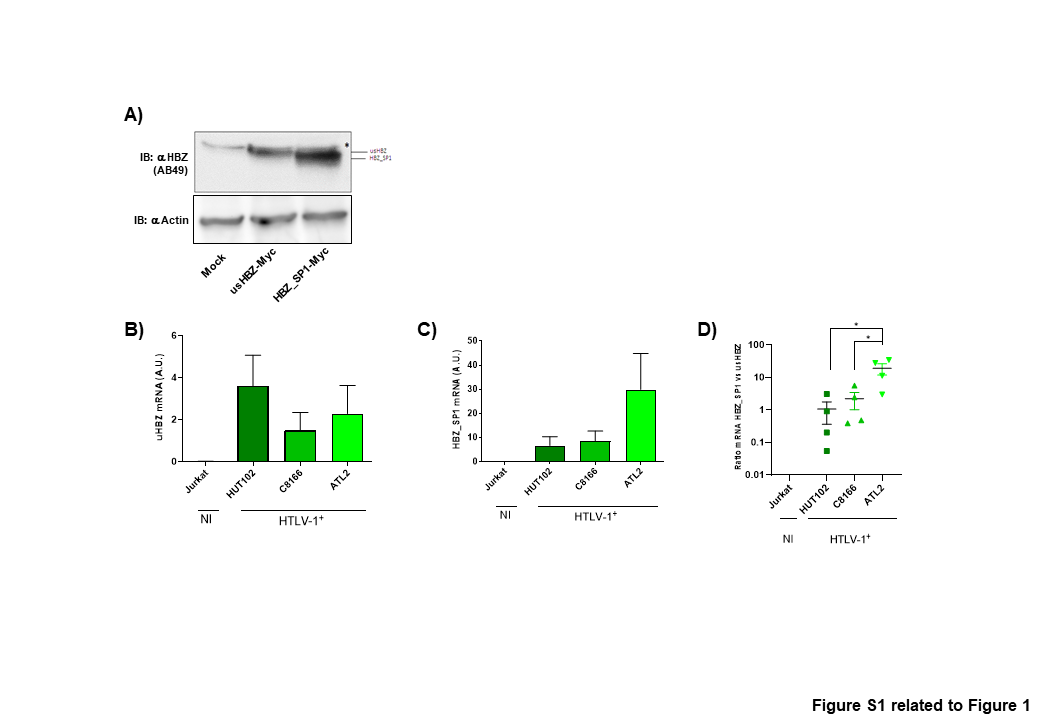

### supplementary Figure 2

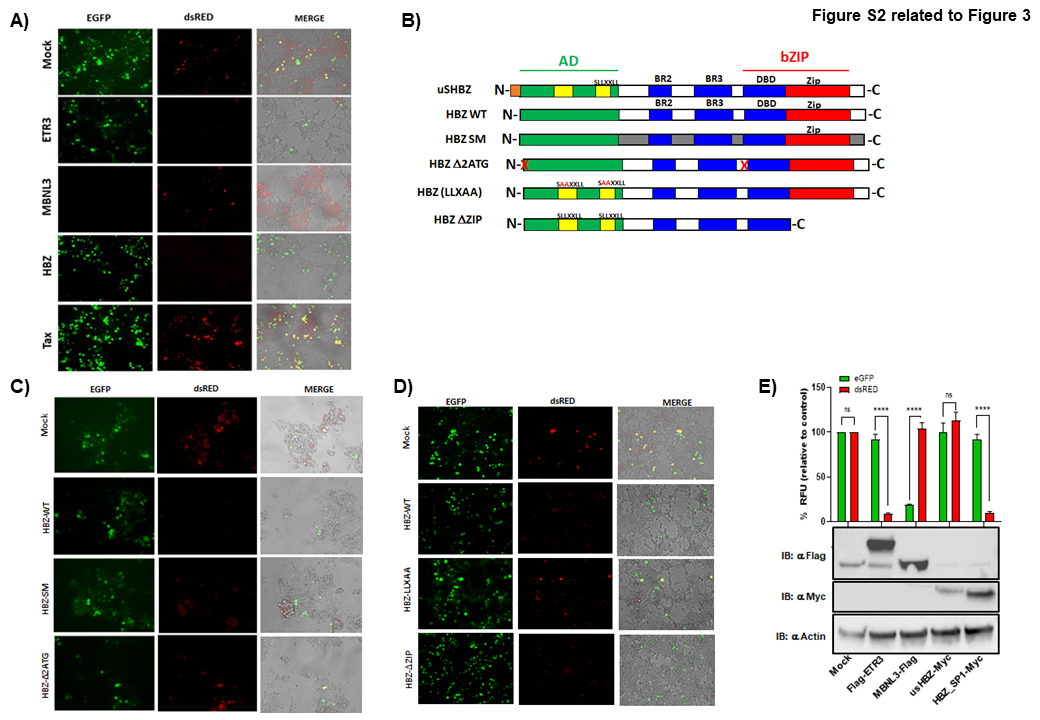

### supplementary Figure 3

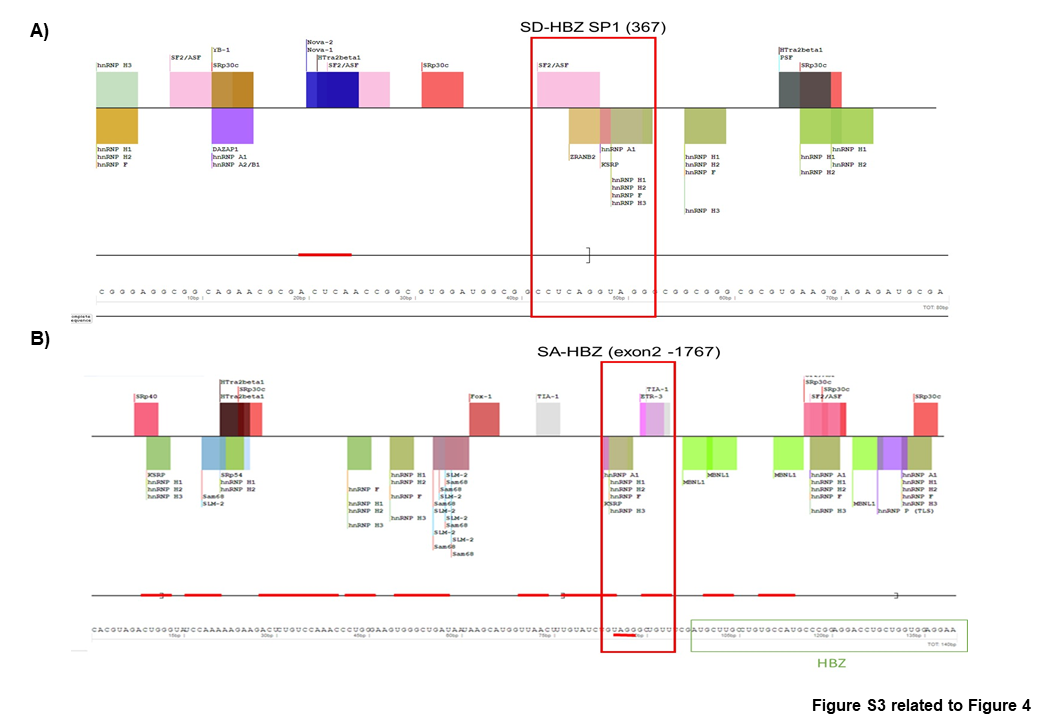

### supplementary Figure 4

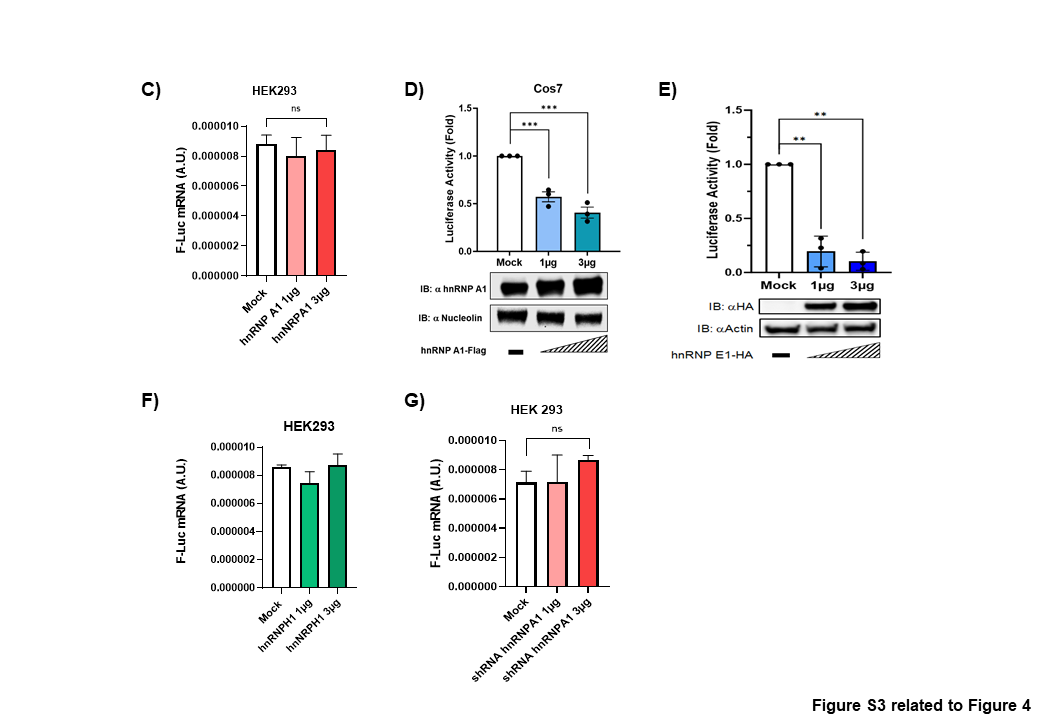

### supplementary Figure 5

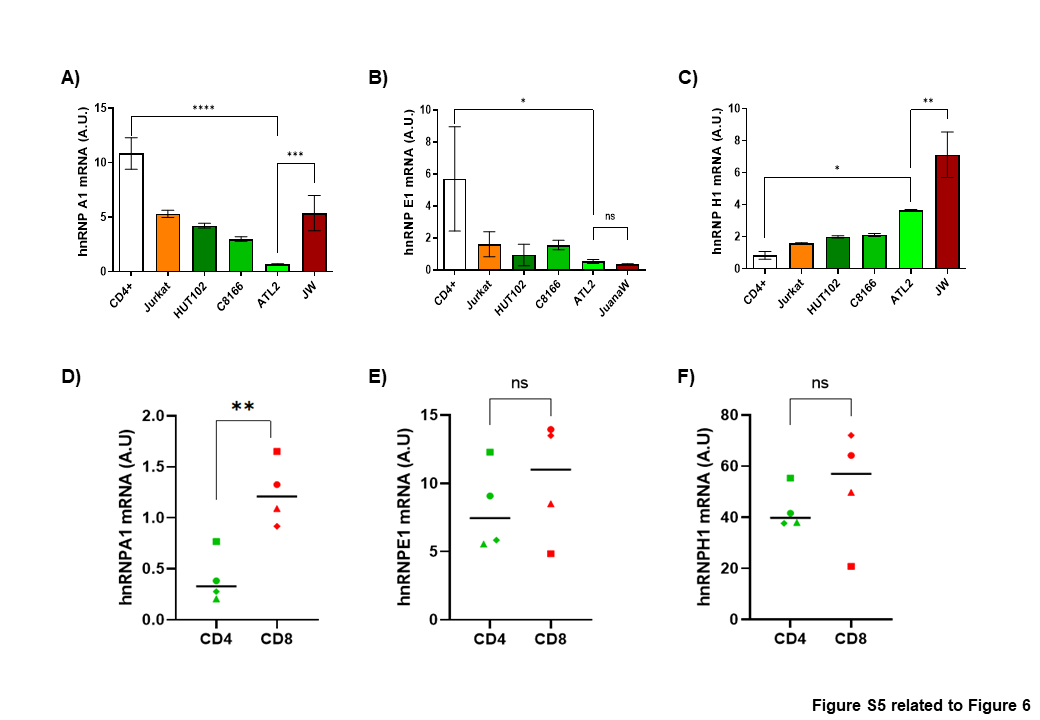
