## supplementary Tables 1 & 2 for "Antagonistic regulation of HBZ splicing by hnRNPA1 and hnRNPH1 drives HTLV-1 leukemogenesis"

**Supplementary Table 1 : List of primers used for RT-qPCR**

| Name | Unique Assay ID: |
| --- | --- |
| hnRNPA1 | qHsaCED0056861 |
| hnRNPC | qHsaCED0005488 |
| hnRNPD | qHsaCID0012561 |
| hnRNPE1/PCBP1 | qHsaCED0056788 |
| hnRNPF | qHsaCED0046661 |
| hnRNPH1 | qHsaCED0046601 |
| hnRNPH2 | qHsaCED0044167 |
| hnRNPK | qHsaCED0046277 |
| hnRNPL | qHsaCID0011499 |
| hnRNPM | qHsaCID0017273 |
| hnRNPR | qHsaCID0009072 |
| hnRNPU | qHsaCID0036652 |
| SF2/ASF (SRSF1) | qHsaCED0036564 |
| SRSF2 | qHsaCED0045938 |
| SRSF3 | qHsaCID0036417 |
| SRSF4 | qHsaCID0007515 |
| SRSF5 | qHsaCED0004829 |
| SRSF6 | qHsaCED0044018 |
| SRSF9 | qHsaCID0012852 |
| ETR3 | qHsaCED0042115 |
| MBNL1 | qHsaCED0037624 |
| RNPS1 | qHsaCED0038600 |
| KSRP | qHsaCID0011555 |
| ZRANB2 | qHsaCID0014684 |
| U2AF1 | qHsaCED0022481 |

| Genes: | sequence | TM |
| --- | --- | --- |
| HPRT -FW | AAGGGCATATCCTACAACAAAC | 60° |
| HPRT -Rev | GGTCAGGCAGTATAATCCAAAG | 60° |
| Tax-FW | ACCAACACCATGGCCCA | 60° |
| Tax-Rev | GAGTCGAGGGATAAGGAAC | 60° |
| HBZ-Fw | ATGGCGGCCTCACCGTCGCAG | 66° |
| HBZ-Rev | GGTCAGGCAGTATAATCCAAAG | 66° |

**Supplementary Table 2 : List of** **primary antibodies**

| Name | type | Compagny |
| --- | --- | --- |
| HBZ (clone AB49) | Mouse Monoclonal | Eurogentec |
| Tax (168B17-46-34) | Mouse Monoclonal | AIDS Research and Reference Reagent Program, NIAID, NIH) |
| hnRNP A1 (4B10), | Mouse monoclonal | Santa Cruz Biotechnology |
| hnRNP C1/C2 (4F4) | Mouse monoclonal | Santa Cruz Biotechnology |
| hnRNP E1 (E-2), | Mouse monoclonal | Santa Cruz Biotechnology |
| hnRNP F/H (1G11) | Mouse monoclonal | Santa Cruz Biotechnology |
| hnRNP K (D-6) | Mouse monoclonal | Santa Cruz Biotechnology |
| hnRNP U (3G6) | Mouse monoclonal | Santa Cruz Biotechnology |
| ETR3 (CUG-BP1/2) (B-1) | Mouse monoclonal | Santa Cruz Biotechnology |
| MBNL1 (3A4) | Mouse monoclonal | Santa Cruz Biotechnology |
| C/EBP alpha (C-18) | Rabbit Poluclonal | Santa Cruz Biotechnology |
| Nucleolin C23 (MS-3) | Mouse Monoclonal | Santa Cruz Biotechnology |
| Anti-Flag-M2 (F1804) | Mouse Monoclonal | Sigma-Aldrich |
| anti-Myc (SAB4700447), | Mouse Monoclonal | Sigma-Aldrich |
| anti-β-actin (A3853) | Mouse Monoclonal | Sigma-Aldrich |
| Anti-Firefly Luciferase antibody (EPR17790) | Rabbit Monoclonal | Abcam |

The mouse monoclonal anti-HBZ antibody (clone AB49) generated in-house (Eurogentec) using an immunizing HBZ peptide selected via the AbDesigner online tool ^52^
